# Structural basis for recognition of RALF peptides by LRX proteins during pollen tube growth

**DOI:** 10.1101/695874

**Authors:** Steven Moussu, Caroline Broyart, Gorka Santos-Fernandez, Sebastian Augustin, Sarah Wehrle, Ueli Grossniklaus, Julia Santiago

## Abstract

Plant reproduction relies on the highly regulated growth of the pollen tube for proper sperm delivery. This process is controlled by secreted RALF signaling peptides, which have been previously shown to be perceived by *Cr*RLK1Ls membrane receptor-kinases and leucine-rich (LRR) extensin proteins (LRXs). Here we demonstrate that RALF peptides are active as folded, disulfide bond-stabilized proteins, which can bind to the LRR domain of LRX proteins with nanomolar affinity. Crystal structures of the LRX-RALF signaling complexes reveal LRX proteins as constitutive dimers. The N-terminal LRR domain containing the RALF binding site is tightly linked to the extensin domain via a cysteine-rich tail. Our biochemical and structural work reveals a complex signaling network by which RALF ligands may instruct different signaling proteins – here *Cr*RLK1Ls and LRXs – through structurally different binding modes to orchestrate cell wall remodeling in rapidly growing pollen tubes.

**Significance:** Plant reproduction relies on proper pollen tube growth to reach the female tissue and release the sperm cells. This process is highly regulated by a family of secreted signaling peptides that are recognized by cell-wall monitoring proteins to enable plant fertilization. Here, we report the crystal structure of the LRX-RALF cell-wall complex and we demonstrate that RALF peptides are active as folded proteins. RALFs are autocrine signaling proteins able to instruct LRX cell-wall modules and *Cr*RKL1L receptors, through structurally different binding modes to coordinate pollen tube integrity.

## Introduction

In flowering plants, sexual reproduction depends on the directional, long-distance growth of pollen tubes that deliver sperm cells to the female gametophytes. The polarized and rapid growth of the pollen tube cell depends on the highly dynamic remodeling of its cell wall. The redundant signaling peptides RALF4 and RALF19 ensure pollen tube integrity and growth by interacting with two distinct protein families, the malectin-domain *Cr*RLK1L membrane receptor kinases (1–3) and the cell-wall monitoring leucine-rich repeat (LRR) extensin proteins (LRXs) (4, 5).

Both *Cr*RLK1Ls and LRXs are required for proper pollen tube growth (4–7), but their relative contribution to specific RALF peptide sensing remains to be investigated. Here, we mechanistically dissect how RALF peptides involved in plant reproduction differentially interact with pollen-expressed LRX proteins and *Cr*RLK1L receptors. Our work highlights that RALFs are folded signaling proteins specifically sensed by LRX proteins that monitor cell wall changes.

## Results

There are 36 RALF peptides in *Arabidopsis* (8), with RALF4/19 being involved in the regulation of pollen tube integrity and growth (1, 4). Mature RALF4/19 peptides are 51 amino acids long and contain four invariant cysteines. We hypothesized that they may be folded proteins, stabilized by intra-molecular disulfide bridges. Hence, for biochemical and structural analyses, we expressed RALF4/19 as thioredoxin A fusion proteins by secreted expression in insect cells. We observed secretion of RALF4/19 into the insect cell medium only when co-expressed with members of the LRX family, but neither as a stand-alone protein nor in the presence of the *Cr*RLK1L ANXUR1 (ANX1) (6, 7) (*SI Appendix*, Fig. S1*A*). Efficient secretion of RALF4/19 required the N-terminal half of LRX8/11, lacking their C-terminal extensin domain (*SI Appendix*, Fig. S1). RALF4/19 formed stable complexes with LRX8_33-400_ (amino-acids 33-400) and LRX11_45-415_ in size-exclusion chromatography (SEC) experiments (*SI Appendix*, Fig. S1 *C* and *D*). The presence of both proteins in the respective complexes was verified by mass spectrometry analysis (*SI Appendix*, Fig. S2). Next, we biochemically mapped the RALF4 binding site to the predicted LRR core of LRX8 and LRX11 N-terminal half (*SI Appendix*, Fig. S1 *B* and *E*).

To quantify the interaction of RALF4 and LRX8, we first dissociated the complex at low pH (see methods) (Fig. 1*A*). The LRX8–RALF4 complex could be fully reconstituted when shifting back to cell wall pH (pH 5.0) (9, 10) (Fig. 1*A*). We titrated the RALF4 protein isolated from insect cells (RALF4_folded_ hereafter) into a solution containing isolated LRX8_49-400_ using isothermal titration calorimetry (ITC), and found that RALF4 tightly binds the LRR core of LRX8 with ∼ 1 nM affinity and a binding stoichiometry (N) of ∼ 1 (Fig. 1*B*). Importantly, a synthetic linear RALF4 peptide binds LRX8 with ∼ 20-fold reduced affinity, suggesting that LRX8 may preferentially sense the RALF4_folded_ signaling protein (Fig. 1*B*). In line with this, a K_*d*_ of ∼ 900 nM has been reported for RALF4_linear_ binding to plant extracted LRX8 using biolayer interferometry (4).

**Fig. 1.**
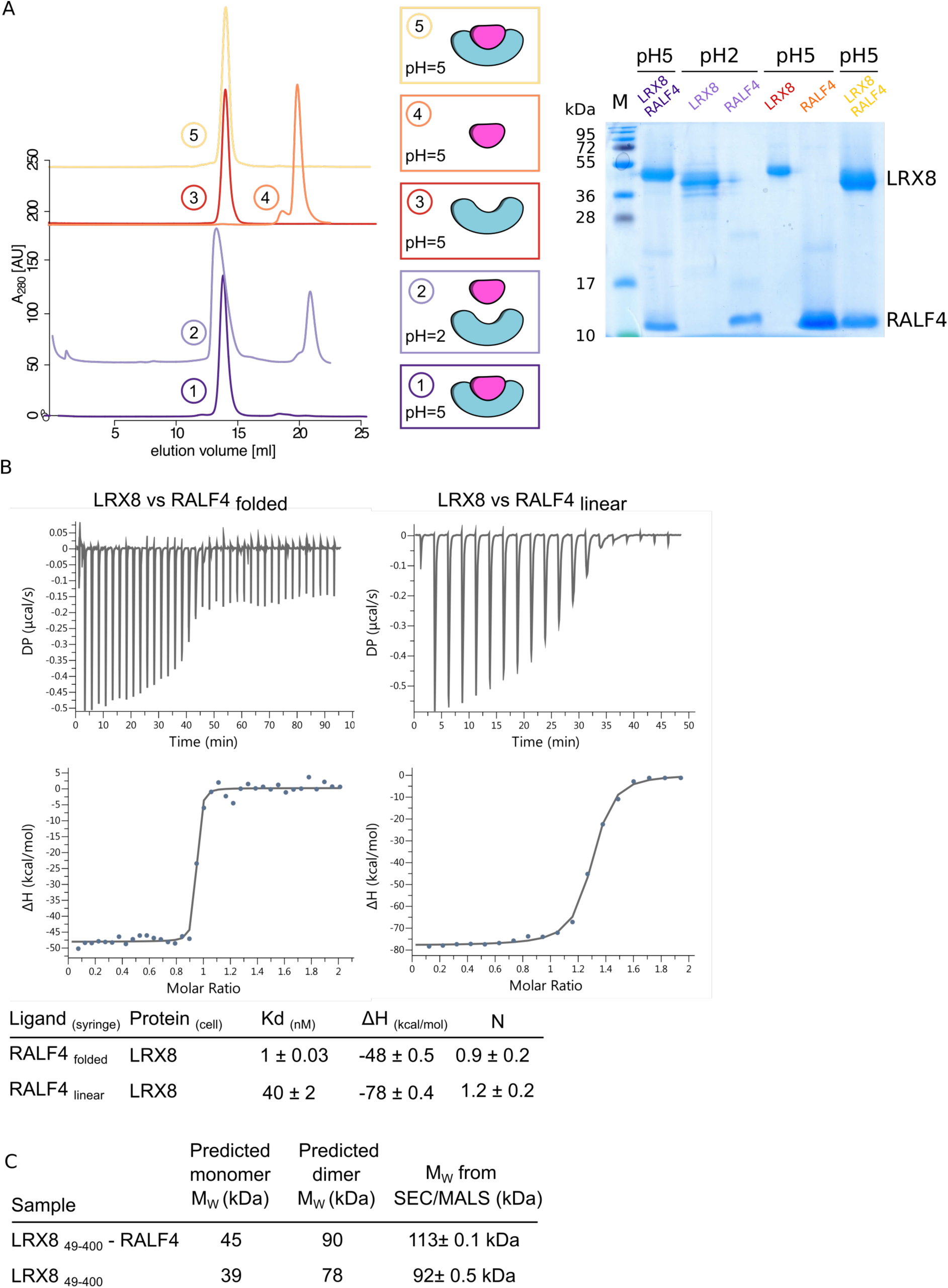
Dimeric LRX proteins bind RALF_folded_ peptides with high affinity. (*A*) Folded RALF4 peptide can be obtained by complex dissociation. LRX8-RALF4 complex purification, dissociation, and reconstitution. Size exclusion chromatography (SEC) graphs of the different steps are plotted on the left panel. From bottom to top: LRX8-RALF4 purified complex at pH 5.0 (dark purple); dissociated LRX8-RALF4 complex at pH 2.0 (light purple). Orange and red runs represent RALF4 and LRX8 peaks, respectively, dialyzed back to pH 5.0. The SEC for the reconstituted complex, using the previously separated proteins, is shown in yellow. Schematics of the different steps, and a SDS gel of the corresponding peaks, are shown alongside. (*B*) Isothermal titration calorimetry (ITC) of LRX8 versus RALF4_folded_ and RALF4_linear_. Raw thermograms of ITC experiments are plotted. The table summarizes the biophysical values obtained. K_*d*_, (dissociation constant) indicates the binding affinity between the two molecules considered (nM). The N indicates the reaction stoichiometry (N=1 for a 1:1 interaction). (*C*) Table of SEC-MALS analysis of apo-LRX8 and LRX8-RALF4 complex. The predicted and measured values are reported in the table for comparison.

Next, we assessed the stoichiometry of LRX–RALF complexes and found LRX8/11 to be constitutive dimers in absence or presence of RALF4_folded_, as judged by analytical SEC and multi-angle light scattering (MALS) experiments (Fig. 1*C* and *SI Appendix*, Fig. S3). The MALS data is consistent with a 2 + 2 complex where each glycosylated LRX protomer binds one RALF4 protein molecule.

The fact that LRX proteins appear to specifically sense RALF_folded_ peptides prompted us to investigate LRX–RALF complex structures. Diffracting crystals were obtained for LRX2–RALF4 and the pollen-specific LRX8–RALF4 complex, determined at 3.2 Å and 3.9 Å resolution, respectively (*SI Appendix*, Table S1). LRX2–RALF4 crystals contain four RALF4-bound dimers in the asymmetric unit, which closely align with each other, and with the LRX8–RALF4 complex (root mean square deviation (r.m.s.d.) is ∼ 0.8 Å comparing 357 pairs of corresponding Cα atoms) (Fig. 2*A*, and *SI Appendix*, Fig. S4). We thus could use the higher resolution LRX2–RALF4 structure to further analyze the signaling complex.

**Fig. 2.**
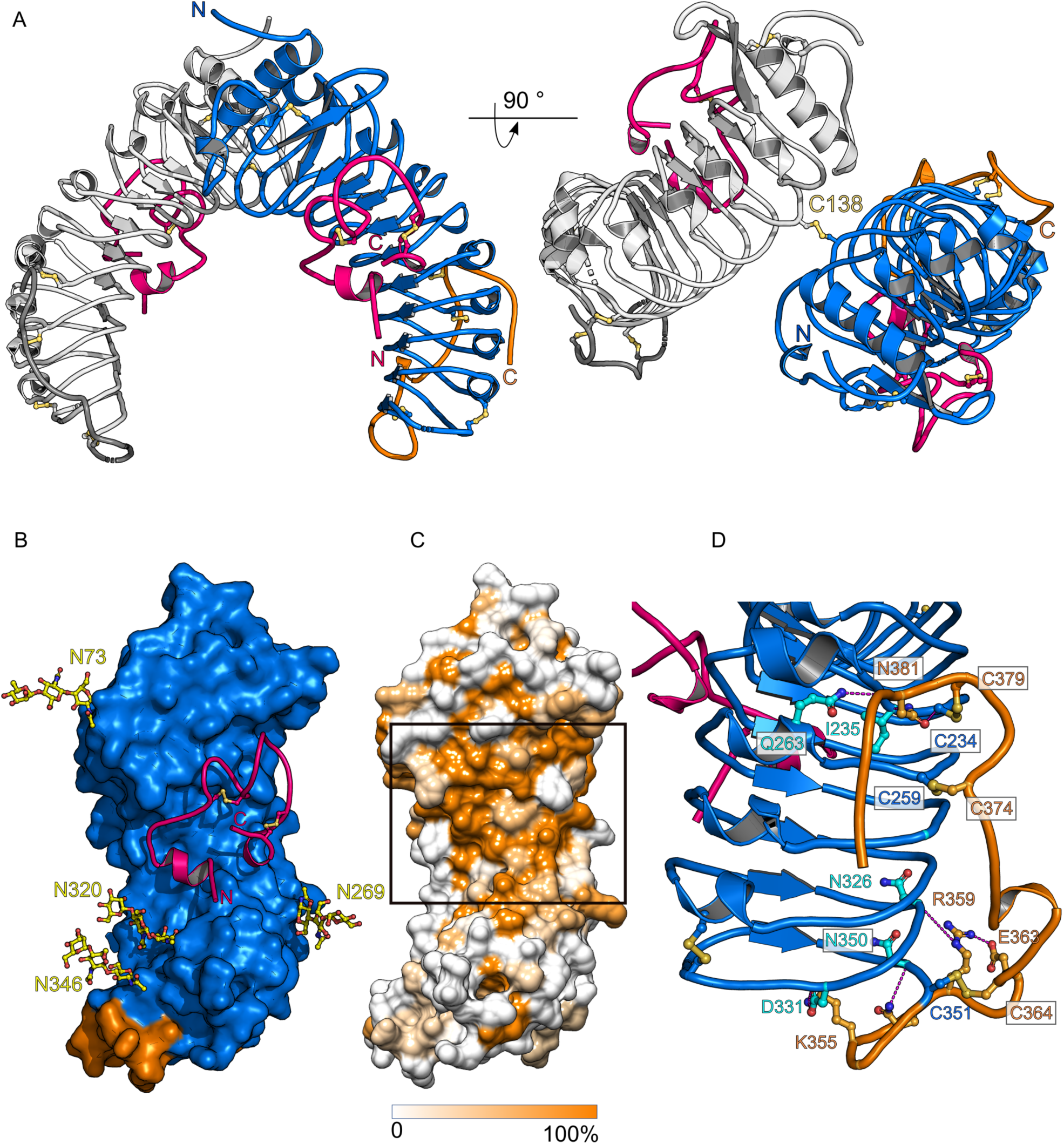
The LRR core of LRX proteins provides a conserved dimer interface, a RALF binding pocket, and a stabilized connection to the extensin domain. (*A*) Front and 90° x-axis rotated view of LRX2 covalently linked homodimer in complex with RALF4 (ribbon diagram). The LRR domain is depicted in blue, the cysteine-rich tail in orange, and the RALF4 peptide is highlighted in pink. The disulfide bridge covalently linking the two LRX protomers is highlighted in yellow. (*B*) Surface view of LRX2 (color code as in **a**) along with cartoon representation of RALF4 peptide (pink), highlighting the LRX RALF binding pocket. N-glycosylations are depicted in yellow. (*C*) Surface representation of LRX2 colored according to the LRX family amino acid conservation. The black rectangle underlines the conservation of the LRX RALF binding pocket. (*D*) Cartoon representation of the C-terminal part of the LRR core and the Cys tail of LRX2 (color code as in *A*). Each connecting disulfide bond is formed between a Cys from the LRR domain (depicted in blue) and the Cys-rich tail (highlighted in orange). Interface residues making polar contacts are shown as sticks and hydrogen bonds are depicted as dotted lines (in magenta).

A structural homology search with the program DALI (11) revealed that the LRX2 protein is closely related to the extracellular domain of known LRR receptor kinases, such as the immune receptor FLS2 (*DALI Z*-score of 33.5, r.m.s.d. of ∼ 1.7 comparing 299 corresponding Cα atoms). LRX2 comprises 11 LRRs sandwiched by canonical N- and C-terminal capping domains, and a cysteine-rich protrusion that represents the N-terminal part of the extensin domain (Fig. 2 *A* and *D*). The LRX2 and LRX8 complex structures confirm that LRX proteins form constitutive dimers, covalently linked by a conserved disulfide bond (Fig. 2*A*, *SI Appendix*, Figs. S4 and S5). One RALF4 peptide is bound to each LRX protomer in the dimer, consistent with our MALS and ITC experiments in solution (Figs. 1 *B* and *C*, and 2*A*). The LRR core provides a binding platform for RALF4_folded_ (Fig. 2*B* and *SI Appendix*, Fig. S4*B*). Despite the moderate resolution of our structures, we located difference electron density accounting for the entire RALF4 folded peptide, again supporting its tight interaction with LRX proteins (*SI Appendix*, Fig. 6*A*). The mature RALF4 protein consists of a short N-terminal alpha-helix, followed by a long loop region that is shaped by two disulfide bonds and a final one-turn 3_10_-helix at the C-terminus. The rather unstructured RALF4 peptide adopts a defined conformation when binding to the LRX LRR domain by making extensive contacts with the binding pocket (Fig. 2*B* and *SI Appendix*, Figs. S4 and S5). The RALF binding pocket is highly conserved among the known LRX family members, suggesting that other LRX proteins may bind different RALF peptides in a similar conformation (Fig. 2*C* and *SI Appendix*, Fig. S5). This particular binding configuration exposes the least conserved regions of the peptide to the solvent (*SI Appendix*, Fig. S7).

In our structures, we also captured the cysteine-rich tail that links the C-terminal LRR capping domain of LRX2 and LRX8 to the cell-wall anchored extensin domain. The cysteine-rich linker adopts similar conformations in the LRX2 and LRX8 structures (*SI Appendix*, Fig. 4*C*). The cysteine-rich tail folds back onto the LRR domain and connects via a ladder of disulfide bridges with LRRs 7-11. Several conserved polar and hydrophobic residues further stabilize this LRR–linker interface (Fig. 2*D* and *SI Appendix*, Fig. S5).

**Fig. 3.**
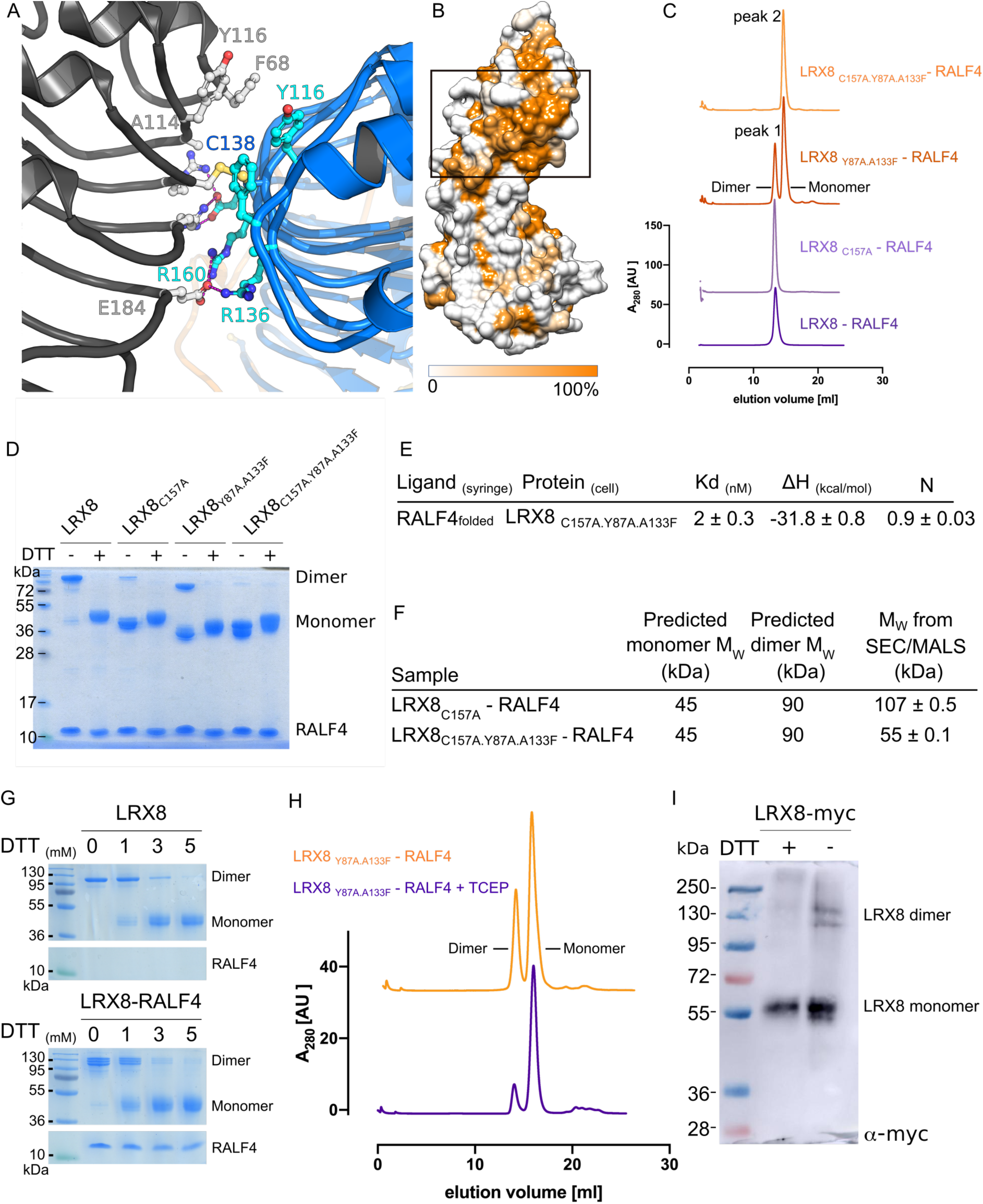
LRX homodimer interface is further stabilized by a central disulfide bridge. (*A*) Close-up view of the dimer interface in LRX2. Details of the interactions between the two protomers (depicted in blue and grey) are highlighted. The Cys (C138) mediating the disulfide bond formation is depicted in yellow. The ionic and hydrogen bonds are shown as dotted lines (in magenta). (*B*) Side surface representation of LRX2 colored according to LRX family sequence conservation. The black rectangle depicts the LRX dimer interface. (*C*) The dimer interface of LRX proteins is stabilized by a central disulfide bridge. SEC runs of wild-type LRX8-RALF4, LRX8_C157A_-RALF4, LRX8_Y87A.A133F_-RALF4, and LRX8_C157A.Y87A.A133F_-RALF4. Peak 1 and 2 correspond to the dimeric and monomeric forms of LRX, respectively. (*D*) SDS-PAGE analysis in the presence/absence of DTT of the mutants shown in *C*. (*E*) Isothermal titration calorimetry (ITC) of LRX8_C157A.Y87A.A133F_ versus RALF4_folded_. The table summarizes the biophysical values obtained. K_*d*_, (dissociation constant) indicates the binding affinity between the two molecules considered (nM). (*F*) SEC-MALS summary table for the LRX8_C157A_-RALF4 and LRX8_C157A.Y87A.A133F_-RALF4 complexes in solution at pH 5.0, along with the predicted molecular weights. (*G*) *In vitro* validation of covalently linked LRX8 homodimers in denaturing conditions. SDS-PAGE gels of LRX8 alone (left) and in complex with RALF4 (right), treated with increasing DTT concentrations (0, 1, 3, and 5 mM). Dimers run above 100kDa while monomers run around 50kDa. (*H*) Shift of the monomer-dimer equilibrium under reducing conditions in solution. LRX8_Y87A.A133F_-RALF4 run in the absence (orange) and presence (purple) of TCEP. (*I*) Detection of monomer and dimer species of transiently expressed LRX8_1-400_ in tobacco leaves, ± DTT.

**Fig. 4.**
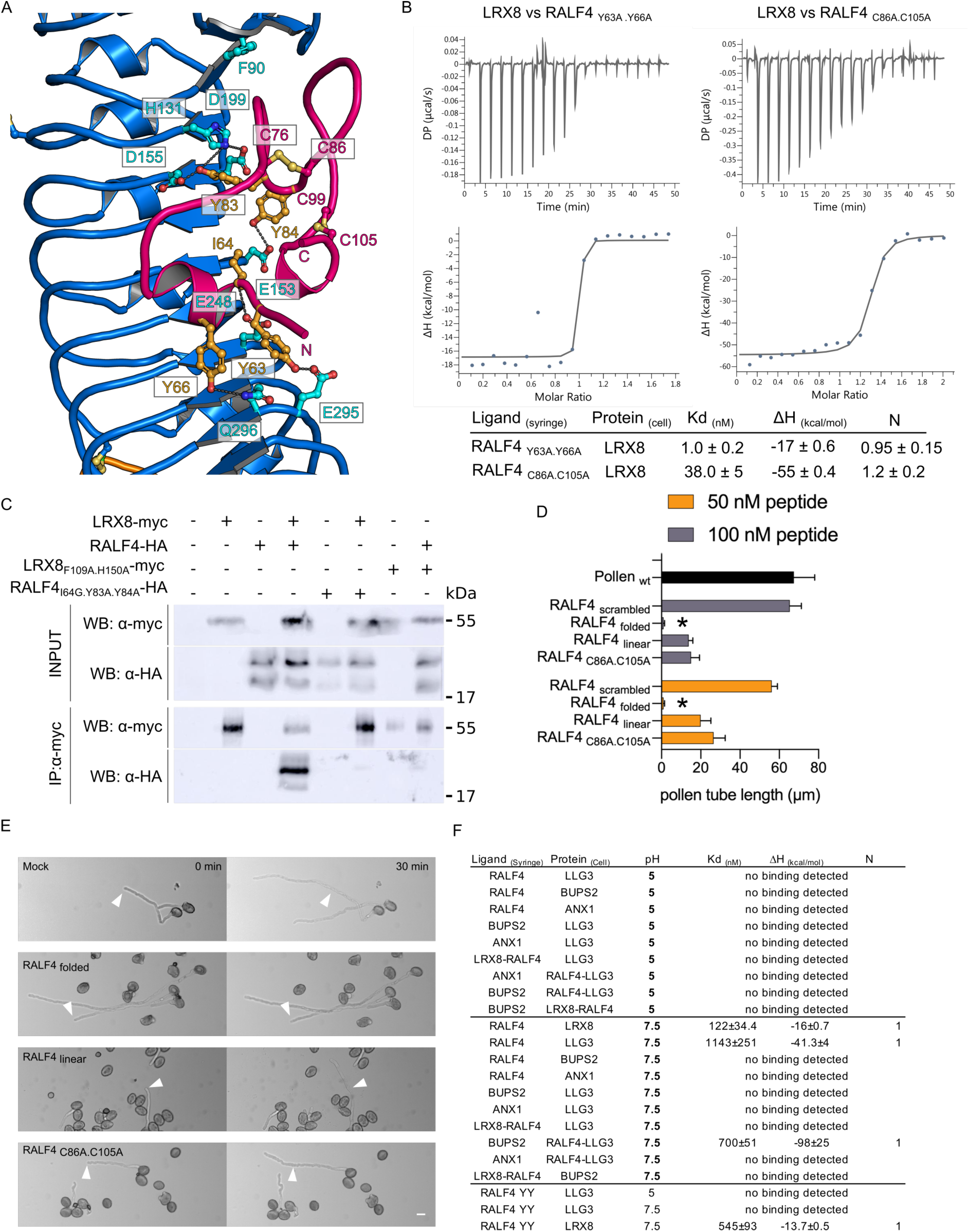
The RALF4 folded conformation is important for its bioactivity. (*A*) Close-up view of the LRX2-RALF4 binding pocket. A large surface of the RALF4 peptide (in pink) directly interacts with the LRR core (in blue) of LRX2. Polar contacts of RALF4 with LRX2 are shown as dotted lines (in grey). The two disulfide bridges responsible for the folding of the peptide, C76-C86 and C99-C105, are depicted as sticks in light orange. (*B*) ITC binding experiments of LRX8 *vs* RALF4_C86A.C105A_ and RALF4_Y63A.Y66A_ mutants. Table summaries for dissociation constant (K_*d*_), binding stoichiometries (N), and thermodynamic parameters are shown alongside. (*C*) *In vivo* validation of the LRX8-RALF4 complex structure. Co-immunoprecipitation (Co-IP) of LRX8-myc and RALF4-HA proteins transiently expressed in tobacco were performed using an anti-myc antibody (IP:α-myc). RALF4-HA peptides were detected with an anti-HA antibody in the IP elution. (*D*) Growth rate of wild-type pollen tubes after the addition of RALF4_folded_, RALF4_linear_, RALF4_scrambled_, and RALF4_C86A.C105A_ peptides at different concentrations after 30 minutes (Data were analyzed by one-way ANOVA followed by Tukey’s test as post hoc, considering p≤0.05 as significantly different; data shown are mean ± s.e.m. of three biological replicates, n=28 each. Distinct letters mean statistically significant differences within groups, ng = no growth after peptide addition). (*E*) Effect of RALF4_folded_, RALF4_linear_, and RALF4_C86A.C105A_ peptides on wild-type pollen tubes growing in semi-solid medium at two time points after adding them to a concentration of 50 nM. Arrowheads indicate the position of the tip of selected pollen tubes at time point 0 min. Scale bar, 10 µm. (*F*) Binding matrix of RALF4 peptide with LRX8 and *Cr*RLK1L/LLGs membrane proteins in acidic and alkaline conditions. ITC table summary with dissociation constants (K_*d*_), binding stoichiometries (N), and thermodynamic parameters (ΔH).

The LRX homodimer is stabilized by a central disulfide bridge (Figs. 2*A* and 3*A*, and *SI Appendix*, Fig. S4*A*), which contributes to a dimer interface formed by highly conserved hydrophobic and polar amino acids (642 Å^2^ buried surface area) (Fig. 3 *A* and *B*, *SI Appendix*, Fig. S5 and Table S2). We probed the LRX8 dimer interface by mutational analysis: LRX8 still behaves as a dimer in solution when the disulfide bond was disrupted by mutating Cys157 (Cys138^LRX2^) to Ala, suggesting that the LRX intermolecular disulfide bond is not essential for structural integrity of the dimer. Simultaneous mutation of Tyr87 (Phe68^LRX2^) to Ala and Ala133 (Ala114^LRX2^) to Phe resulted in a monomer-dimer equilibrium, which we could quantitatively shift to a LRX8 monomer by additionally introducing the Cys157Ala mutation (Fig. 3 *C* and *F*, *SI Appendix*, Fig. S8 and Table S2). Both dimeric and monomeric LRX8 forms retain the capacity to bind RALF4, with the monomer binding the folded peptide with wild-type affinity (∼2 nM) (Fig. 3 *D*-*F*). To assess the role of the central cysteine in a changing redox environment, we incubated purified LRX8 and LRX8–RALF4 complex with increasing concentrations of the reducing agent DTT in denaturing conditions. In both cases, the dimer could be converted into monomers (Fig. 3*G*). In addition, we could also quantitatively shift the oligomeric equilibrium of LRX8_Y87A.A133F_ towards the monomeric species in the presence of a reducing agent in SEC experiments (Fig. 3*H*). Consistent with this, we could detect both oligomeric species of a myc-tagged LRX8_1-400_, when transiently expressed in tobacco leaves, in the absence of DTT (Fig. 3*I*).

We next dissected the RALF4 binding mode to LRX8. Interaction between the N-terminal alpha-helix (residues 64 – 69) of RALF4_folded_ with the LRR core of LRXs are mediated by Tyr63, Ile64, and Tyr66, making hydrogen bonding and hydrophobic interactions with LRX8 residues Gln314 (Gln296^LRX2^), Phe221(Phe202^LRX2^), Glu266 (Glu248^LRX2^), and Glu313 (Glu295^LRX2^) (Fig. 4*A*). A RALF4 Tyr63Ala/Tyr66Ala double mutant still bound LRX8 with wild-type affinity, suggesting that the N-terminal helix in RALF peptides is not a major determinant for high-affinity binding to LRX proteins (Fig. 4*B*). However, mutation of the Cys86 and Cys105 to Ala, targeting the two disulfide bonds observed in our RALF4 crystal structures, led to a ∼ 20-fold reduction in binding when compared to wild-type RALF4_folded_. This suggests that the structural stabilization of the RALF4_folded_ loop region (residues 70 –101) by the Cys76–Cys86 and Cys99–Cys105 disulfide bonds is required for high affinity binding to LRX8 (Fig. 4 *A* and *B*), in good agreement with the reduced binding affinity observed for the linear RALF4 peptide (Fig. 1*B*). We thus mutated the LRX–RALF4 loop interface and tested if stable LRX8–RALF4 complexes could still form. Co-expression of LRX8 Phe109Ala/His150Ala (Phe90^LRX2^/His131^LRX2^) and wild-type RALF4 led to a reduced expression of the LRX8 protein in insect cells. Secretion of the RALF4 peptide was not detected, suggesting that the mutations in the LRX RALF binding interface inhibit the formation of a stable LRX8–RALF4 complex. The same behavior was observed when mutating the corresponding residues in the RALF4 loop region (Ile64Gly/Tyr83Ala/Tyr84Ala) (*SI Appendix*, Fig. S9). Consistently, the LRX8 and RALF4 mutant proteins also failed to form stable complexes *in planta* as judged by co-immunoprecipitation experiments in tobacco leaf extracts (Fig. 4*C*). Altogether, our experiments define the disulfide bond-stabilized loop in RALF4_folded_ as a major determinant for LRX binding.

Based on these findings, we compared the bioactivity of linear, folded, and cysteine-mutated RALF4 signaling peptides in pollen tube growth assays (4). The inhibitory effect of RALF4 peptides on pollen tube elongation was quantified using two peptide concentrations: 50 and 100 nM (Fig. 4*D*). Addition of a sequence-permutated version of RALF4 (RALF4_scrambled_) had no effect on pollen tube growth, whereas RALF4_folded_ completely inhibited growth at both concentrations. The linear and cysteine-mutated RALF4 peptides could only partially suppress pollen tube growth (Fig. 4 *D* and *E*), in good agreement with our structural and quantitative biochemical data (Figs. 1*B*, 2*A* and 4*B*).

It has previously been reported that *Cr*RLK1Ls FERONIA (FER) and THESEUS1 (12, 13) are receptors for RALF peptides with binding affinities in the nM to µM range (1, 14–16). We thus tested if RALF4_folded_ can directly bind to ANX1 and BUDDHA’S PAPER SEAL2 (BUPS2), which have been genetically implicated in pollen tube growth (1, 6, 7). We expressed the ectodomains of ANX1 and BUPS2 by secreted expression in insect cells as previously reported (2), and quantified their interaction with RALF4_folded_ in ITC assays in acidic and alkaline conditions (16, 17) (Fig. 4 *F*). We found no detectable interaction between RALF4_folded_ and either ANX1 or BUPS2 at pH 5.0 or pH 7.5, respectively. Like FER, the GPI-anchored protein LORELEI, expressed in the female gametophyte, is required for fertilization (18, 19), and LORELEI-LIKE proteins (LLGs) co-immunoprecipitate with FER in the presence of RALF peptides (20). We thus produced the ectodomain of the pollen-expressed LLG3 (*SI Appendix*, Fig. S10 and S11) and found that it binds RALF4_folded_ with a K_*d*_ of ∼1 µM at pH 7.5, with no detectable binding at pH 5. There was also no detectable interaction between LLG3 and either ANX1 or BUPS2 in the absence of the peptide. However, BUPS2, but not ANX1, only binds a LLG3-RALF4_folded_ complex with a dissociation constant of ∼ 700nM at pH 7.5 (Fig. 4 *F*), suggesting specific recognition between *Cr*RLK1Ls and LLG-RALFs in the pollen. The pH also modulates LRX8-RALF4 binding, with affinity decreasing ∼100 fold between pH 5 and pH 7.5 (Fig. 1*B* and 4*F*). Strikingly, the Tyr63Ala/Tyr66Ala mutant in the RALF4 N-terminal alpha-helix, which still binds LRX8 with wild-type affinity at both pHs (5 and 7.5), disrupts the interaction with LLG3 at pH 7.5 (Fig. 4 *B* and *F*). This suggests that LLG/*Cr*RLK1Ls and LRX proteins may both sense RALF peptides in pollen tubes; however, with vastly different binding affinities and mechanistically distinct peptide binding modes. Despite being seemingly involved in the same process, we could not observe any biochemical interaction between LRX proteins and *Cr*RLK1Ls/LLGs in the presence or absence of RALF4 (Fig. 4 *F*).

## Discussion

Taken together, our mechanistic analysis of RALF signaling peptides in pollen tube growth reveal that RALFs are folded, disulfide bond-stabilized signaling proteins, which interact with the N-terminal LRR domain of LRX proteins in the cell wall with high affinity. A recent independent NMR study on refolded RALF8 produced in bacteria, also confirms the presence of two disulfide bonds in the peptide and reports that the peptide lacks a specific folded state by itself in solution (21). It is of note that RALF4, when bound to LRX, adopts a defined conformation exposing a highly basic surface patch, which could mediate targeted interactions with other proteins or cell wall components (*SI Appendix*, Fig. S6*B*). Binding of RALFs to the LRR region could thus affect the overall conformation of the LRX extensin domain, which our structures reveal to be tightly linked to the LRR core. Moreover, since polarized growth is sensitive to changes in the cell wall redox environment (22, 23), the oligomeric state of LRX proteins may be affected by changes in the redox state of the plant cell wall. Such redox-controlled oligomerization of LRX proteins may add an additional layer in the modulation of RALF signaling. In this respect, it is noteworthy that plant cell wall thioredoxins have been identified as putative LRX interactors (24). That we do not observe direct interaction between LRX and *Cr*RLK1Ls/LLGs signaling modules, could be explained by the autocrine signaling mode of RALF4/19 during pollen tube growth (1, 4). On the one hand, RALF4/19 signal via the ANX1/2 *Cr*RLK1Ls to affect processes inside the pollen tube through the cytoplasmic receptor-like kinase MARIS (4, 25); on the other hand, RALF4/19 may signal to control processes outside the cell through their interaction with LRXs in the cell wall. This is consistent with the finding that, unlike mutants with reduced RALF4/19 activity (4), mutants lacking multiple pollen-expressed LRXs cannot be suppressed by dominant active MARIS (*SI Appendix*, Fig. S12) (26), indicating that RALF4/19 work in different but converging pathways to fine-tune pollen tube integrity. Our binding data between RALFs, LRXs and *Cr*RLK1Ls/LLGs, together with previous studies (16), suggest that this autocrine signaling could be additionally regulated by cell-wall pH fluctuations to modulate cell wall integrity (9). Altogether, these findings suggest that RALFs have the capacity to instruct different signaling proteins to coordinate cell wall remodeling during pollen tube growth (*SI Appendix*, Fig. S13). Our structural, quantitative biochemical, and physiological experiments all support that folded rather than linear RALF peptides represent the bioactive ligands for LRX and *Cr*RLK1L/LLG proteins. This point should be considered in future studies aimed at dissecting the contributions of LRXs and *Cr*RLK1Ls to the diverse RALF-mediated signaling processes in plants.

## Acknowledgements

We thank V. Olieric for providing beam-time and the staff at beam line PXIII of the Swiss Light Source (SLS), Villigen, for technical assistance during data collection.

## Data and materials availability

All data is available in the main text or in the supplementary materials.

## Materials and Methods

### Protein expression and purification

Codon optimized synthetic genes for expression in *Spodoptera frugiperda* (Invitrogen GeneArt, Germany), coding for *Arabidopsis thaliana* LRX2 (residues 1-385), LRX8 (residues 33-400, 49-400, and 49-373), LRX11 (residues 45-415, 64-415, and 64-388) domains were cloned into a modified pFastBac (Geneva Biotech) vector, providing a TEV (tobacco etch virus protease) cleavable C-terminal StrepII-9xHis tag. RALF4 (residues 58-110) and RALF19 (residues 58-110) mature peptide sequences were N-terminally fused to TRX A (Thioredoxin A) in a pFastBac vector driven by a 30K signal peptide (28). For protein expression, *Trichoplusia ni* Tnao38 cells (29) were co-infected with a combination of LRX and RALF virus with a multiplicity of infection (MOI) of 3 and incubated for 1 day at 28°C and two days at 22°C at 110 rpm. The secreted complexes were purified from the supernatant by sequential Ni^2+^ (HisTrap excel; GE Healthcare; equilibrated in 25 mM KP_i_ pH 7.8, 500 mM NaCl) and StrepII (Strep-Tactin Superflow high capacity; IBA; equilibrated in 25 mM Tris pH 8.0, 250 mM NaCl, 1 mM EDTA) affinity chromatography. Proteins were further purified by size-exclusion chromatography on a Superdex 200 increase 10/300 GL column (GE Healthcare), equilibrated in 20 mM sodium citrate pH 5.0, 150 mM NaCl. For crystallization and biochemical experiments, proteins were concentrated using Amicon Ultra concentrators (Millipore, MWCO 3,000 and 30,000). Proteins were analyzed for purity and structural integrity by SDS-PAGE and mass spectrometry.

### Mass spectrometry

For LC-MS/MS analysis of LRX8-RALF4, 5 µL of complex (1 mg/ml) were diluted in 20 µL of the following buffer: 50 mM ammonium bicarbonate, 10 mM TCEP (triscarboxyethylphosphine) and 40 mM chloroacetamide. Samples were incubated 45 min in the dark at RT, and then digested with 0.1 µg of sequencing-grade trypsin or chymotrypsin (Promega). Samples were incubated at 37 °C for 4 hours, and digestion reaction stopped with 2µl of 10% formic acid. They were then diluted 10x with loading buffer (2% acetonitrile, 0.05% trifluoroacetic acid) and injected on a Fusion Tribrid orbitrap mass spectrometer (Thermo Fisher Scientific) interfaced to a Dionex RSLC 3000 nano-HPLC. Peptides were separated on a 65 min. gradient from 4% to 76% acetonitrile in 0.1% formic acid at 0.25 µL/min on an in-home packed C18 column (75 μm ID x 40 cm, 1.8 μm, Reprosil Pur, Dr. Maisch). Full MS survey scans were performed at 120’000 resolution. In data-dependent acquisition controlled by Xcalibur 4.1 software (Thermo Fisher), a top speed precursor selection strategy was applied to maximize acquisition of peptide tandem MS spectra with a maximum cycle time of 1.5 s. HCD (normalized collision energy: 32%) or EThcD (supplemental HCD activation energy: 25%) fragmentation mode were used with a precursor isolation window of 1.6 m/z. MS/MS spectra were acquired at a 15’000 resolution, and peptides selected for MS/MS were excluded from further fragmentation during 60s. For data analysis, tandem mass spectra were searched using PEAKS software (version 8.5, Bioinformatics Solutions Inc., Waterloo, Canada) against a custom database containing common contaminants and the sequences of the proteins of interest. Mass tolerances used were 10 ppm for the precursors and 0.02 Da for collision-induced dissociation fragments. Activities of proteases considered were semi-trypsin or semi-chymotrypsin (one specific cut) with 2 missed cleavages. Carbamidomethylation of cysteine was specified as a fixed modification. N-terminal acetylation of protein and oxidation of methionine were specified as variable modifications. PEAKS results were imported into the software Scaffold 4.8.5 (Proteome Software Inc., Portland, OR, USA) for validation of MS/MS based peptide (minimum 90% probability) and protein (min 95 % probability) identifications, dataset alignment as well as parsimony analysis to discriminate homologous hits.

For LC-MS/MS analysis of LRX11-RALF4, 10 µL of complex (10 mg/ml) were diluted in 20 µL of the following buffer: 8M Urea, 50 mM TEAB (triethylammonium bicarbonate), 5mM TCEP (triscarboxyethylphosphine) and 20 mM chloroacetamide. After 1h incubation at RT, the sample was then diluted adding 90 µL of 50mM TEAB, split into two aliquots of 55 ul and digested with 0.2 µg of trypsin or chymotrypsin. Samples were incubated at 37 °C for 2.5 hours, and digestion reaction stopped with 1µl of 10% formic acid and 200 µL of loading buffer. They were injected on a Fusion Tribrid orbitrap mass spectrometer interfaced to a Dionex RSLC 3000 nano-HPLC, as above. Full MS survey scans were performed at 120’000 resolution. In data-dependent acquisition controlled by Xcalibur 4.1 software, a top speed precursor selection strategy was applied to maximize acquisition of peptide tandem MS spectra with a maximum cycle time of 0.6 s. HCD (normalized collision energy: 32%) fragmentation mode was used with a precursor isolation window of 1.6 m/z. MS/MS spectra were acquired in the linear trap and peptides selected for MS/MS were excluded from further fragmentation during 60s. For data analysis, tandem mass spectra were searched using Mascot (Matrix Science, London, UK; version 2.6.2) against a custom database containing common contaminants and the sequences of the proteins of interest. Mass tolerances used were 10 ppm for the precursors and 0.5 Da for collision-induced dissociation fragments. Activities of proteases considered were semi-trypsin (one specific cut) or chymotrypsin, with 2 or 3 missed cleavages, respectively. Carbamidomethylation of cysteine was specified as a fixed modification, and oxidation of methionine as variable modification. Mascot results were imported into the software Scaffold for validation of MS/MS based peptide (minimum 95% probability) and protein (min 95 % probability) identifications, dataset alignment as well as parsimony analysis to discriminate homologous hits.

### Multi-angle light-scattering (MALS)

MALS was used to assess the monodispersity and molecular weight of the apo LRXs and in complex with RALF4. Samples containing 100 µg of protein were injected into a Superdex 200 increase 300/10 GL column (GE Healthcare) using a HPLC system (Ultimate 3000, Thermo Scientific) at a flow rate of 0.5 ml.min^−1^, coupled in-line to a multi-angel light scattering device (miniDAWN TREOS, Wyatt). Static light-scattering was recorded from three different scattering angles. The scatter data were analyzed by ASTRA software (version 6.1, Wyatt). Experiments were done in duplicate with SD indicated in the figures.

### Protein complex dissociation

LRX-RALF complexes were obtained *in vivo* by co-expression in insect cells. The apo proteins were obtained by complex dissociation *in vitro* using a low pH buffer. After purification of the LRX-RALF complex (see above), the protein complex was incubated for 1 h at 4°C in the following buffer: 20 mM citric acid pH 2, 150 mM NaCl. Next, the dissociated proteins were separated by gel filtration, using a Superdex 200 increase 10/300 GL column (GE Healthcare) equilibrated in the same buffer. Each protein was then collected separately and dialyzed against 20 mM citric acid pH 5, 150 mM NaCl. To test whether proteins were still folded and functional, proteins were again run in a Superdex 200 increase 10/300 GL column (GE Healthcare) in 20mM citric acid pH 5, 150mM NaCl. We also tested, for each batch of dissociated protein, that the complex would be fully reconstituted when mixing the proteins in equimolar proportions. Additionally, we tested the *in vivo* activity in pollen tube growth assays of the dissociated peptide (folded) versus a synthetic linear version of RALF4 of the same sequence.

### Isothermal titration calorimetry (ITC)

Experiments were performed at 25°C using a MicroCal PEAQ-ITC (Malvern Instruments, UK) with a 200 µL standard cell and a 40 μL titration syringe. LRX8 protein and RALF peptides were gel-filtrated into either pH 5 ITC buffer (20 mM sodium citrate pH 5.0, 150 mM NaCl) or pH 7.5 ITC buffer (20 mM HEPES pH 7.5, 150 mM NaCl). A typical experiment consisted of injecting 2 μl of a 50 μM solution of the RALF synthetic peptides (RALF4 _linear_ and RALF4 _C86A.C105A,_ sequence and manufacturer below) into 5 μM LRX8 (49-400) solution in the cell at 150 s intervals. For the higher affinity experiments between RALF4_folded_ protein and LRX8, the injection volume was reduced to 1 μl. The Experiments using RALF, *Cr*RLK1Ls, and LLGs were typically performed with 10µM protein in the cell and 100µM ligand in the syringe, and an injection pattern of 2 μl. ITC data were corrected for the heat of dilution by subtracting the mixing enthalpies for titrant solution injections into protein free ITC buffer. Experiments were done in duplicates and data was analyzed using the MicroCal PEAQ-ITC Analysis Software provided by the manufacturer.

### Crystallization and data collection

Crystals of the LRX2_29-385_-RALF4 and LRX8_33-400_–RALF4 complexes developed at room-temperature in hanging drops composed of 1.0 μL of protein solution (20 mg/mL) and 1.0 μl of the following crystallization buffers, respectively : 20% [w/v] PEG 3,350, 0.1 M Bis-Tris pH 5 and 0.2 M sodium acetate and 17.5% [w/v] PEG 8,000, 0.1 M Bis-Tris pH 7 and 0.2 M sodium citrate. Drops were suspended above 0.6 mL of crystallization buffer. For data collection, LRX2-RALF4 crystals were transferred into crystallization buffer supplemented with 15 % (v/v) ethylene glycol and snap frozen in liquid nitrogen. In the case of the LRX8-RALF4 complex, crystals were snap frozen in the presence of 15% glycerol as cryo protectant. A 3.2 Å and 3.9 Å dataset native was collected at beam-line PXIII of the Swiss Light Source (SLS), Villigen, CH. Data processing and scaling was done in XDS (v. June 2017) (30).

### Structure determination and refinement

The structure of the LRX2–RALF4 complex (PDB-ID: 6QXP) was determined by molecular replacement using the program PHASER, implemented in the Phenix Mrage pipeline (31). Search models were selected using the program HHPRED (32) and tested iteratively in the Mrage pipeline. A solution using a fragment of the LRR ectodomain of the plant immune receptor FLS2 (PDB-ID 4MNA) yielded a solution with eight molecules in the asymmetric unit. The search model, which showed 28% sequence identity with LRX2 target was further improved using the program CHAINSAW in CCP4(33, 34). The final solution was used for NCS averaging and density modification in the program PHENIX.RESOLVE (35). The resulting map was readily interpretable and the LRX2-RALF4 complex structure was build and completed by iterative round of manual model building in COOT (36) and restrained TLS in phenix.refine. The structure of LRX8-RALF4 (PDB-ID: 6QWN) was solved by molecular replacement in PHASER (37) and using the LRX2-RALF4 as search model. Inspection of the final models with phenix.molprobity (38) reveal good stereochemistry (Table 1). Diagrams were prepared with PYMOL (https://pymol.org/) or CHIMERA (39).

### Transient protein expression in tobacco leaves

*Nicotinia benthamiana* plants were grown for three weeks prior to agroinfiltration. Appropriate *Agrobacterium tumefaciens* cultures were grown in YEB media at 28°C until reaching an O.D of 0.6. Cultures were centrifuged 10 min at 4000 r.p.m. Cells were resuspended in fresh infiltration media (50 mM MES, 2 mM NaH_2_PO4, 0,5% (m/v) saccharose, 100 µM acetosyringone, pH 5.6) and let on the shaker for 2 h in the dark. For co-infiltration of two *Agrobacterium* strains, infiltration media with the appropriate *Agrobacterium* strains harboring the desired constructs were mixed in a 1:1 v/v ratio. Infiltration was performed with a syringe. Plants were grown for 2 more days prior to material collection. Leaves were cut and flash frozen in liquid nitrogen.

### *In vitro* redox sensitivity experiments

To determine the importance of disulfide bridge formation in the LRX homodimers, samples were incubated at 95°C for 5 min in Laemmli buffers containing DTT 0, 1, 3, 5, or 20 mM, respectively. They were subsequently run on denaturing SDS gels for further size identification. In solution, reducing assays were performed by incubating the LRX8_Y87A.A133F_ protein with and without 5 mM TCEP for 1 hour at RT. Samples were then run by size-exclusion chromatography on a Superdex 200 increase 10/300 GL column (GE Healthcare), equilibrated in 20 mM sodium citrate pH 5.0, 150 mM NaCl, in the presence or absence of TCEP.

### Protein extraction, co-immunoprecipitation and Western blot analysis

Transiently expressed LRX8 and RALF4 proteins in tobacco were extracted as follow: 300 mg of plant material were crushed in liquid nitrogen and resuspended in 600 µL of extraction buffer (150 mM Tris pH7.5; 150 mM NaCl; 10% (v/v) glycerol; 10 mM EDTA, 0.5% IGEPAL, cOmplete protease inhibitor (Roche)). Samples were centrifuged at 15000 r.p.m. for 20 min, and the supernatant was then centrifugated twice more to discard residual pellet debris. 50 µL of input sample were kept at this stage, the rest was incubated for 1 h at 4°C with anti-myc or anti-HA magnetic beads (µMACS, Milteny Biotech). Immunoprecipitation was performed according to manufacturer’s specifications, using the extraction buffer for column washes. Standard Laemmli buffer (40) was added to the input and immunoprecipitated samples prior to gel loading. Finally, Western blot analysis was performed with horse radish peroxidase-coupled anti-myc and anti-HA antibodies (MACS, Milteny Biotech) at 1:2000 dilution. Detection was performed using WesternBright™ Sirius (Advansta). Images were taken with a LAS500 BlotImager.

### Pollen germination *in vitro* assays

Open flowers were incubated at 22°C for 30 min in moist incubation boxes. Then, pollen was bound to xilane-coated slides containing germination medium [0.01% boric acid (w/v), 5 mM CaCl2, 5 mM KCl, 1 mM MgSO4, 10% sucrose, pH 7.5]. Pollen grains were pre-incubated in moist incubation boxes for 30-45 min at 30°C and then transferred to 22°C from 30 min to 2h as indicated. After 2 h of *in vitro* pollen germination, peptides were added to pollen germination medium to a concentration of 50 nM and 100 nM. The effect on pollen tube growth was imaged using a Leica DM6000 microscope and analyzed using the ImageJ 1.40g software (rsb.info.nih.gov/ij). RALF4_linear_, RALF4_scrambled_ and RALF4 _C86A.C105A_ synthetic peptides were obtained from PHTD Peptides Industrial Co, Ltd. (Zhengzhou City, China). The rest of the peptides were expressed and produced in insect cells complexed with LRX8 and dissociated as indicated above.

<u>Sequence of RALF4_linear_:</u> RRYIGYDALKKNNVPCSRRGRSYYDCKKRRRNNPYRRGCSAITHCYRYAR

<u>Sequence of RALF4_scrambled_:</u> PTYNSCRRKCKRDRGYAARRYKRYRYVNADIKRNHSGYPCRICSRLYGRN

<u>Sequence of RALF4 _C86A.C105A_</u> RRYIGYDALKKNNVPCSRRGRSYYDAKKRRRNNPYRRGCSAITHAYRYAR

### Generation of transgenic lines

All transgenic lines were generated using the floral dip method (41) with the *Agrobacterium tumefaciens* strain GV3101. The construct pLAT52:MRIR240C-YFP, described previously (25), was used for *Arabidopsis thaliana* transformation. The construct’s expression in pollen tubes was checked and imaged using a Leica DM6000 microscope.

