## Supplemental information for "Structural basis for recognition of RALF peptides by LRX proteins during pollen tube growth"

**This PDF file includes:**

Figs. S1 to S13
Table S1 and S2
References citations for Materials and Methods and SI (27-41)

**Figure S1. RALF4/19 form a tight complex with the LRR core of pollen LRX proteins.**

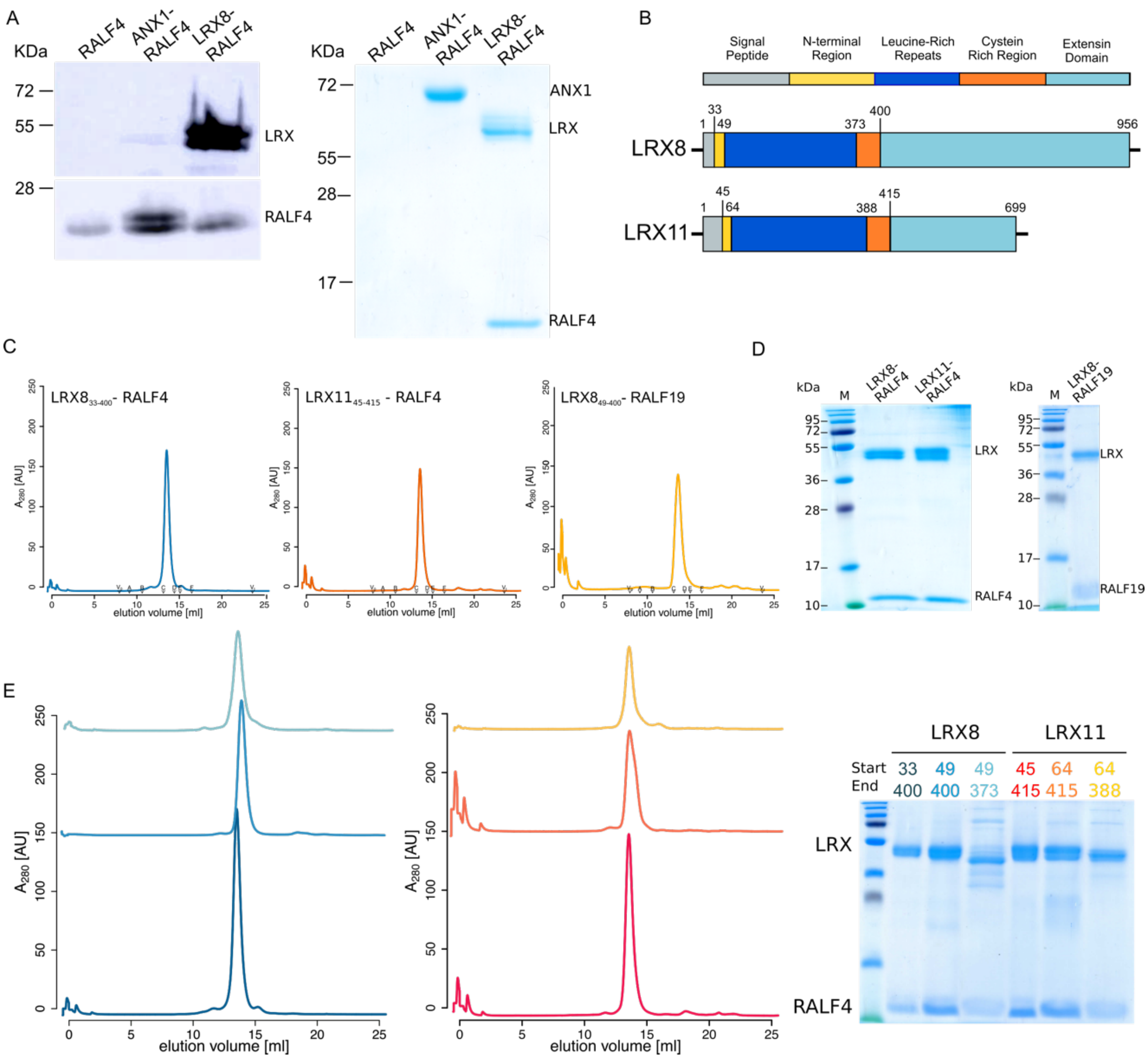

(*A*) Anti-HIS Western blot (left) of insect-cell culture pellets expressing or co-expressing RALF4, ANX1-RALF4, and LRX8-RALF4; SDS-PAGE of corresponding secreted protein fractions (right). (*B*) Schematic overview of LRX8 and LRX11 domains, including the amino acid coordinates according to structural data. (*C*) Analytical size exclusion chromatography (SEC) of LRX8$_{33\text{-}400}$-RALF4, LRX8$_{45\text{-}415}$-RALF4, and LRX8$_{49\text{-}400}$-RALF19 complexes. (*D*) SDS-PAGE of the different peaks corresponding to the SEC experiments of panel (*C*). (*E*) Mapping of the LRX8/LRX11 domain interacting with RALF4 according to the schematic diagram in (*B*). SEC (left) of the LRX8/11-RALF4 complexes. SDS-PAGE (right) of the different SEC peaks. Colors of the SEC plots and the corresponding peak legends are matching.

**Figure S2. LRX8-RALF4 and LRX11-RALF4 mass spectrometry sequence determination.**

A

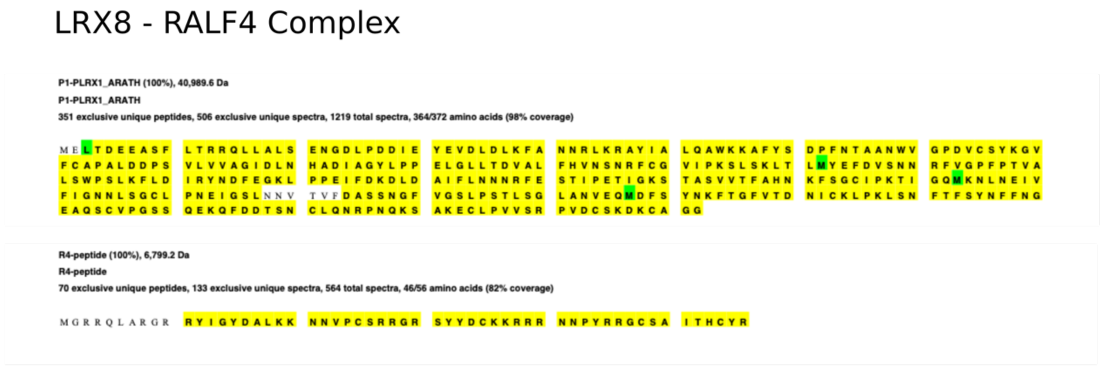

B

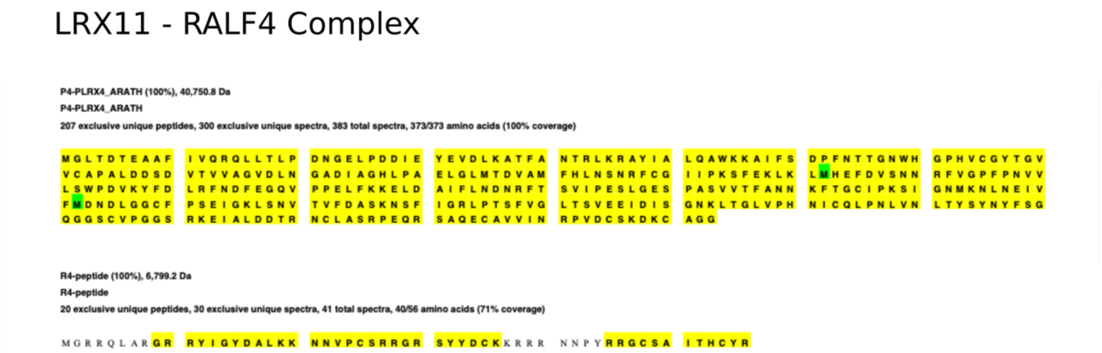

Results from LC-MS/MS analysis of LRX8-RALF4 and LRX11-RALF4 protein complexes. Number of peptides and coverage for each component of the complex is highlighted in yellow. Oxidised methionines are highlighted in green. Leu 3 of LRX8 was identified as the N-terminal, N-acetylated residue, although the modification was only sporadically identified and was thus partial. Cys residues were reduced and alkylated (carbamidomethyl, not shown) and searched as fixed modification. All the sequence coverages are the combined results of tryptic and chymotryptic digestions and separate LC-MS/MS analyses on a high resolution orbitrap Fusion instrument. Samples in (*A*) were analysed by both HCD and EThcD fragmentation, while for results in (*B*) only HCD fragmentation was used. For more details see the Materials and Methods section.

**Figure S3. LRX8 and 11 are constitutive dimers in their apo form or in complex with RALF4.**

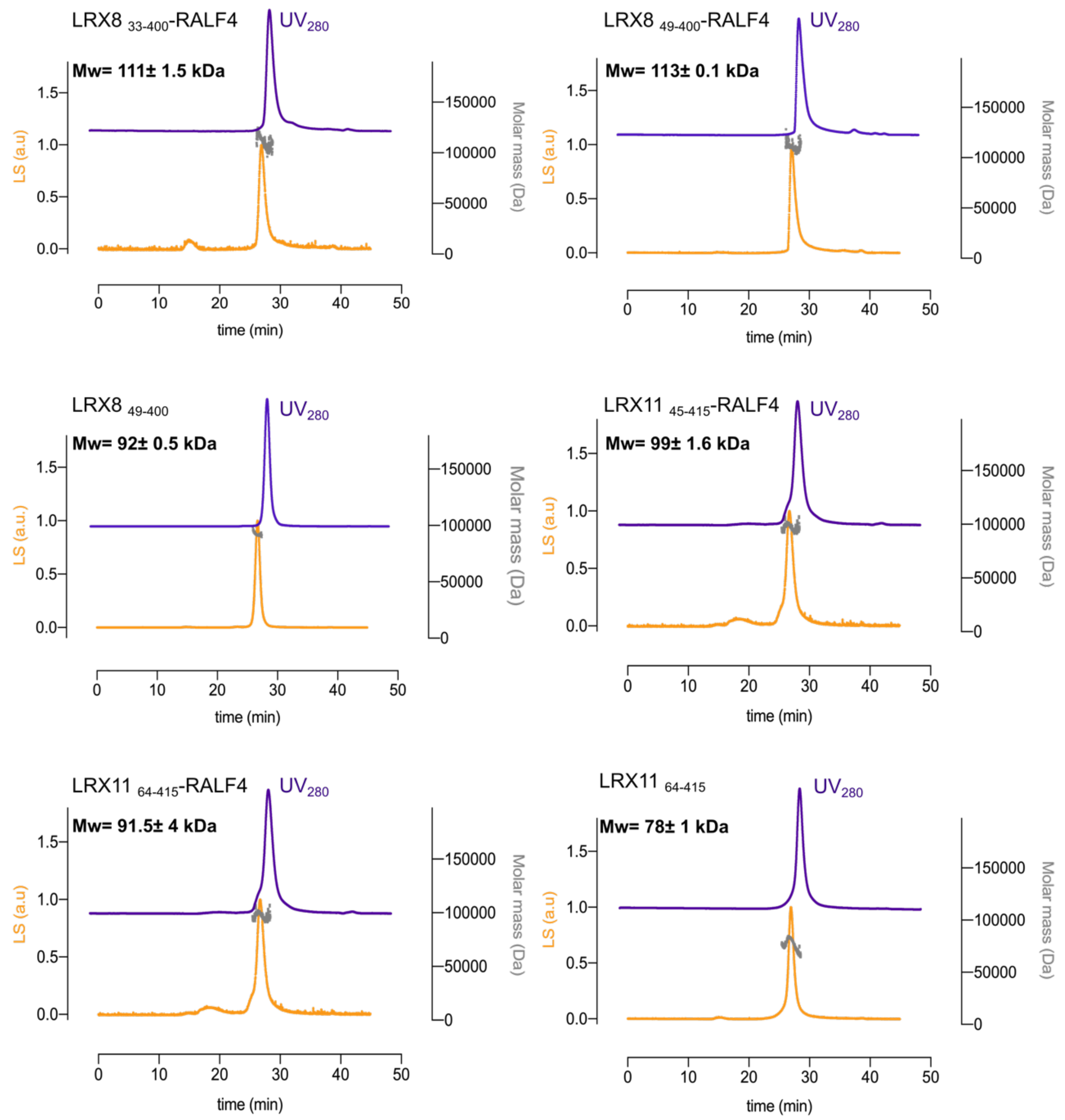

Molecular weight determination of the respective complexes using Multi Angle Light Scattering (MALS). $UV_{280}$ absorption is plotted in purple, light scattering (LS) in orange, and the determined molecular weight (Da) in grey. The molecular weight indicated on each plot is the mean ± SD of two independent measurements.

**Figure S4. LRX8-RALF4 and LRX2-RALF4 complexes share a common architecture.**

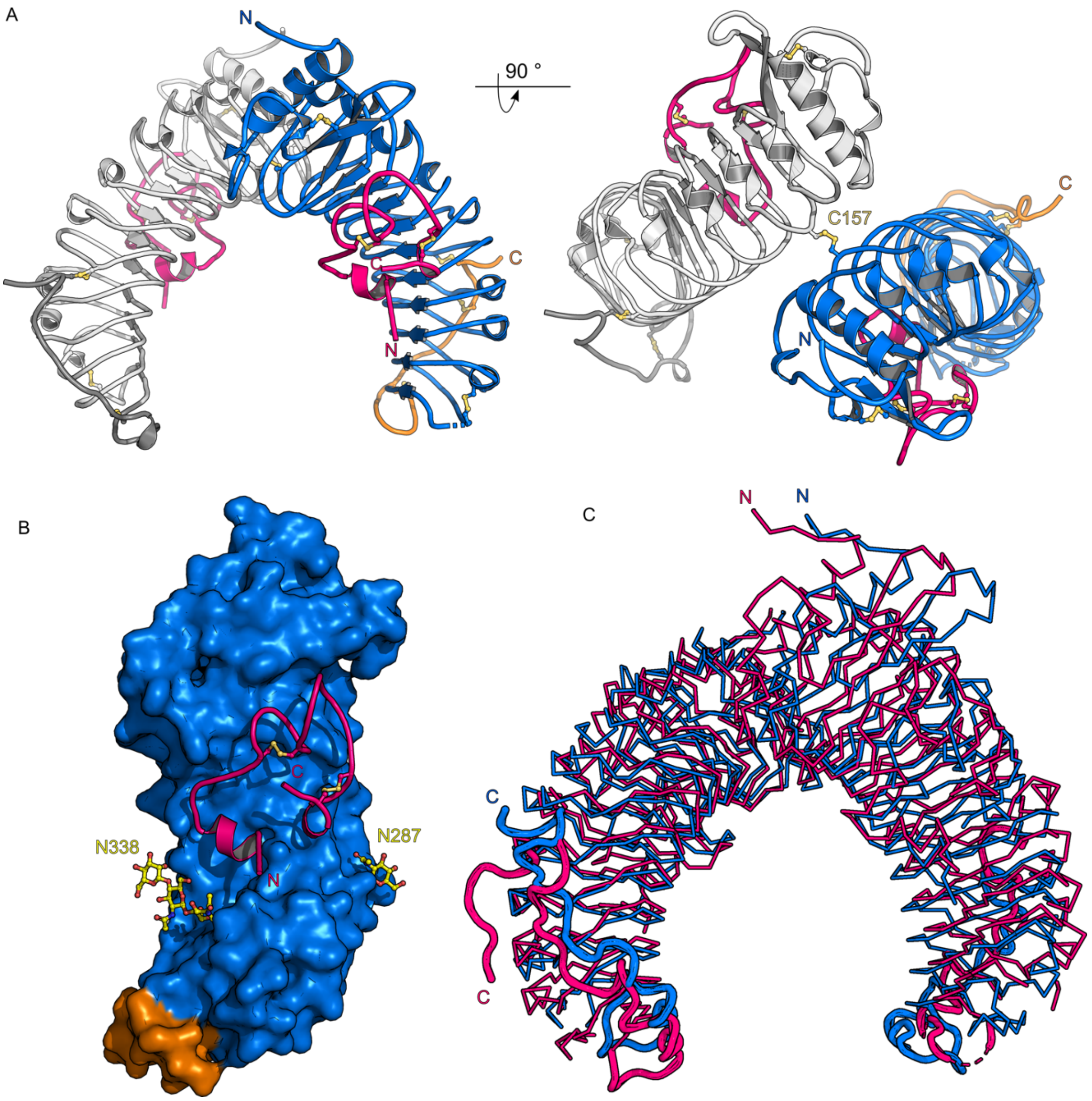

(*A*) Front and 90º x-axis rotated view of LRX8 covalently linked homodimer in complex with RALF4 (ribbon diagram). The LRR domain is depicted in blue, the Cys-rich tail in orange, and the RALF4 peptide is highlighted in pink. The disulfide bridge covalently linking the two LRX protomers is highlighted in yellow. (*B*) Surface view of LRX8 (color code as in A) along with cartoon representation of RALF4 peptide (in pink), highlighting the binding surface of the peptide. N-glycosylations are depicted in yellow. (*C*) Structural superimposition of the LRX2-RALF4 (Cα trace in pink) and LRX8-RALF4 (in blue) complexes. r.m.s.d. of ~ 0.8 Å comparing 357 pairs of corresponding Cα atoms between LRX2-RALF4 and LRX8-RALF4 complexes.

**Figure S5. The LRX2/8-RALF4 complex interfaces are conserved among the LRX protein family.**

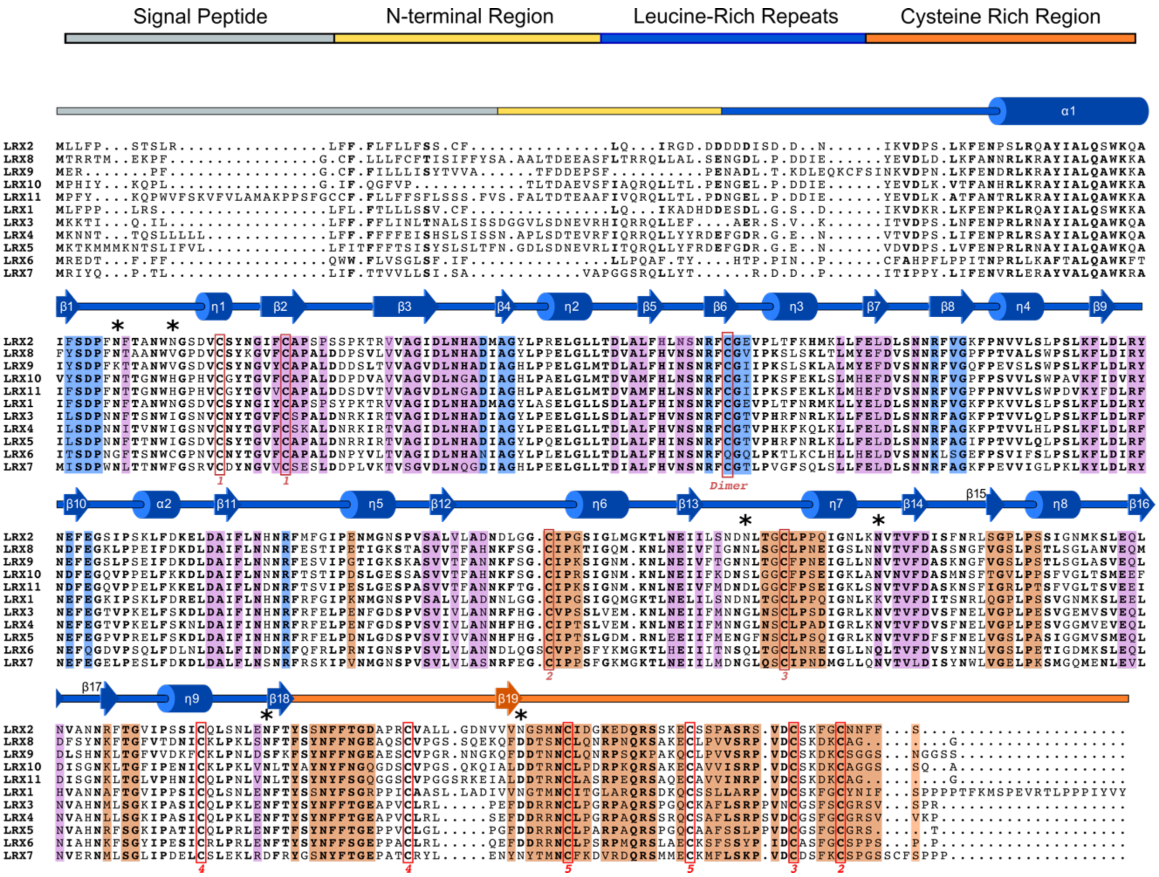

Structure-based sequence alignment of the LRX family members. The alignment includes a secondary structure assignment calculated with the program DSSP (27) and colored according to Fig. 2 and Fig. S1*B*. Cys residues engaged in disulfide bonds are squared in red, with matching pairs indicated by numbers. Predicted and experimentally verified N-glycosylation sites are depicted by black stars. The various interaction surfaces in the LRX2/8-RALF4 complex structures are highlighted according to the following color code: LRX dimer interface in blue, LRX-RALF4 binding pocket in pink, and the interaction surface of the LRX Cys extension with the LRR core of the protein is highlighted in orange.

**Figure S6. RALF4 omit map and surface electrostatic potential of the peptide exposed surface.**

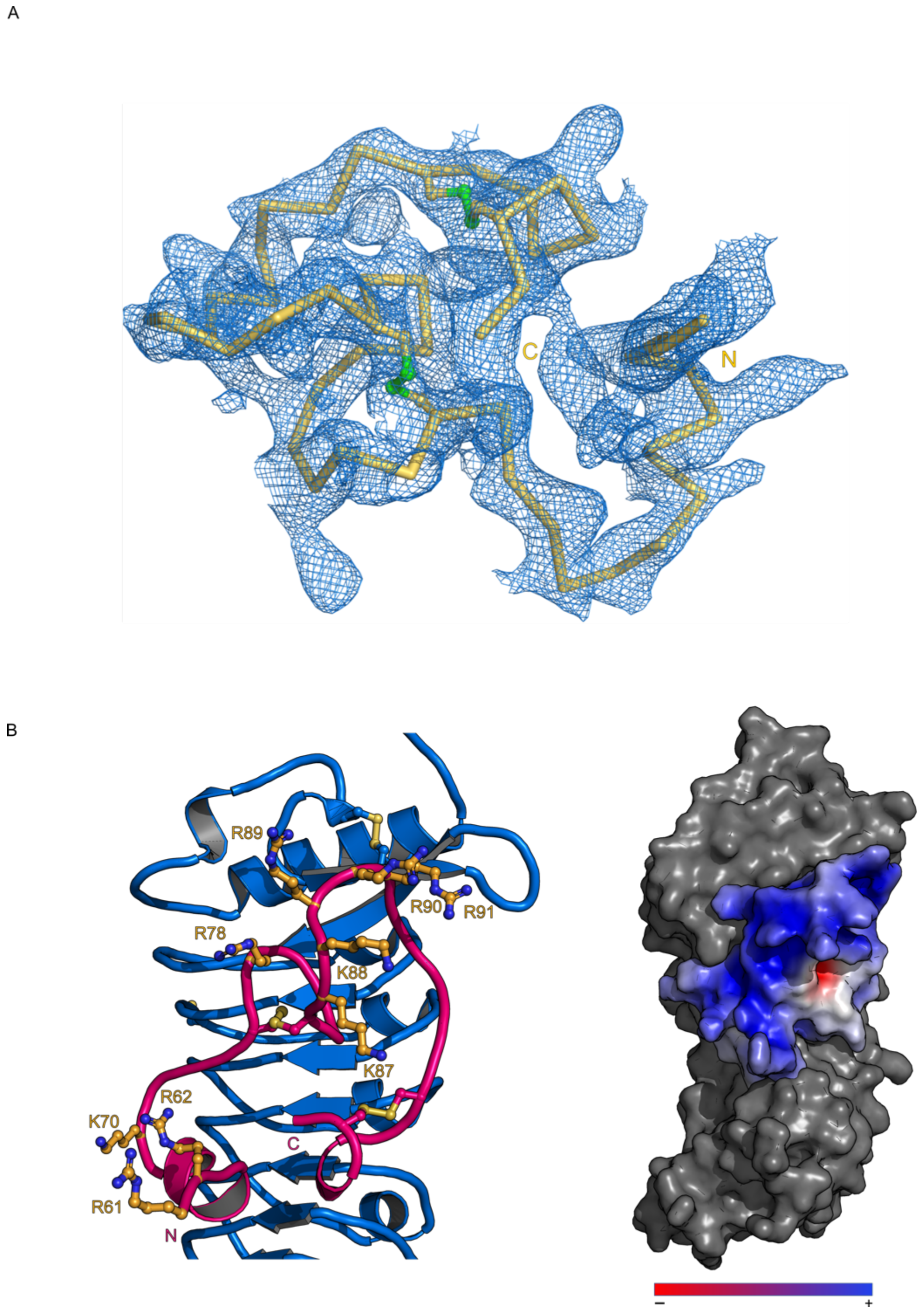

(*A*) RALF4 simulated annealing 2Fo-Fc omit electron density map contoured at 1.1σ. (*B*) RALF4 reveals a positively-charged surface when bound to LRX. Details of the basic nature of RALF4 exposed surface (left). Arg (R) and Lys (K) free residues are shown as sticks in light orange. Electrostatic surface representation of RALF4 when in complex with LRX2 (right). LRX2 is shown in grey and RALF4 solvent-accessible surface electrostatic potential has been calculated using APBS plugin (PyMOL). The potential is given with the negative (red) and positive (blue) contour levels in the range from -8.0 to +8.0 kBT, respectively.

**Figure S7. RALF4 binding interface to LRX is conserved among closely related RALF family members.**

A

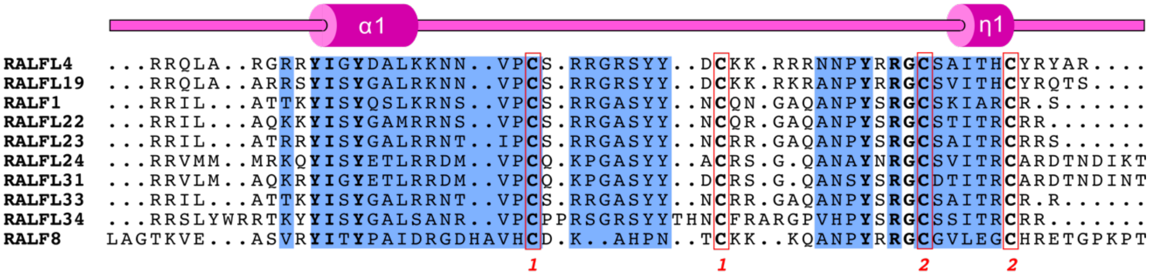

B

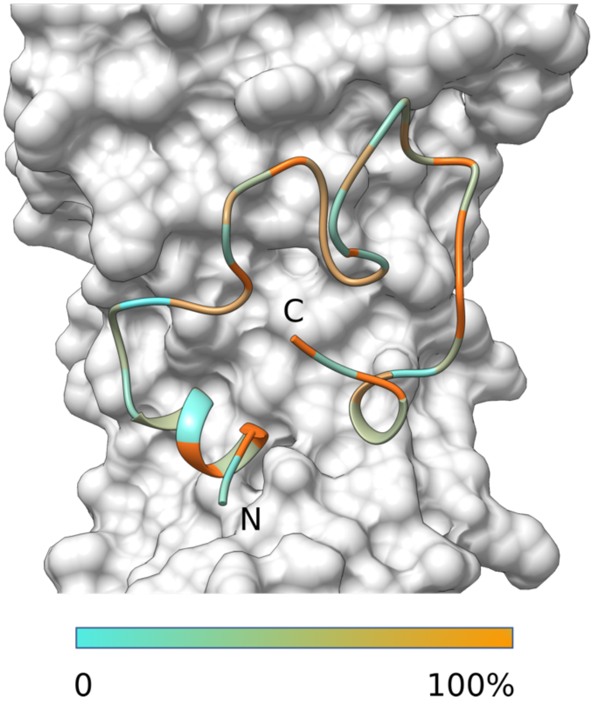

(*A*) Structure-based sequence alignment of RALF4 family related members. The alignment includes a secondary structure assignment calculated with the program DSSP and colored according to Fig. 2. Cys residues engaged in disulfide bonds are squared in red, with matching pairs indicated by numbers. The RALF4-LRX2 interaction surface is highlighted in blue. (*B*) The RALF4 regions directly interacting with the LRX binding pocket are conserved among RALF4-related family members. RALF4 ribbon representation colored according to sequence conservation among RALF peptides indicated in the alignment above.

**Figure S8. Quantitative SEC-MALS analysis of the LRX-RALF dimer interface mutants.**

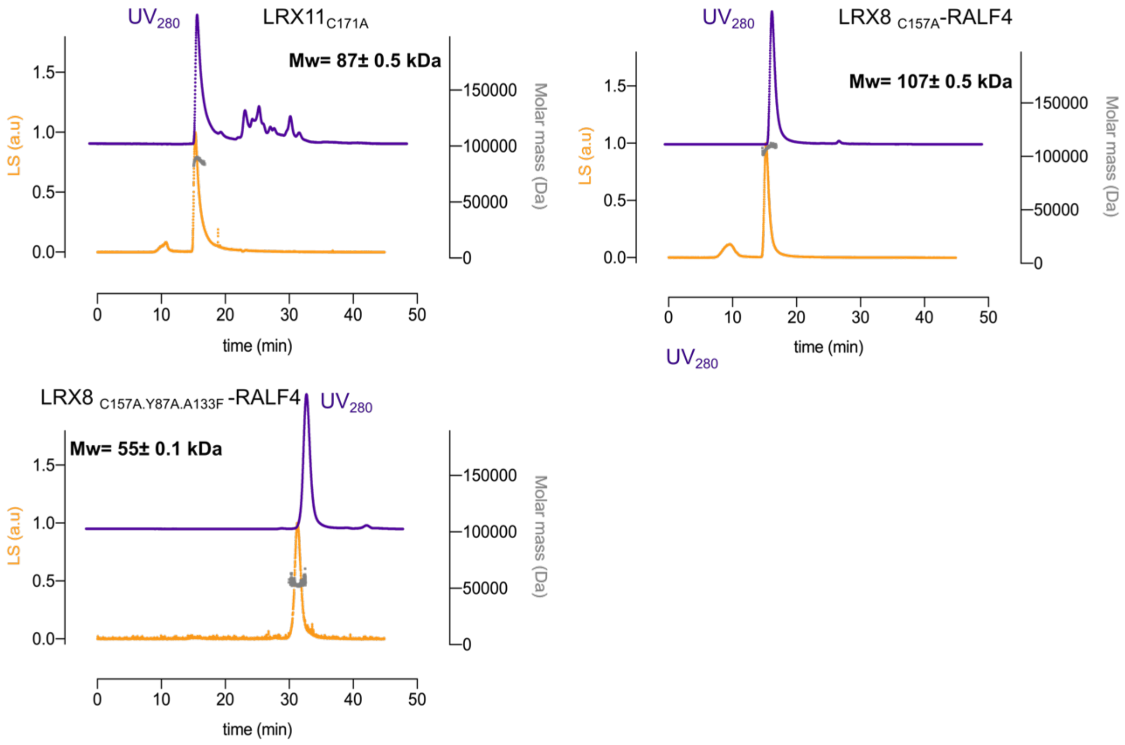

Molecular weight determination of the respective complexes using MALS. $UV_{280}$ absorption is plotted in purple, light scattering (LS) in orange, and the determined molecular weight (Da) in grey. The molecular weight indicated on each plot is the mean ± SD of two independent measurements.

**Figure S9.** ***In vitro*** **validation of the LRX8-RALF4 structure complex.**

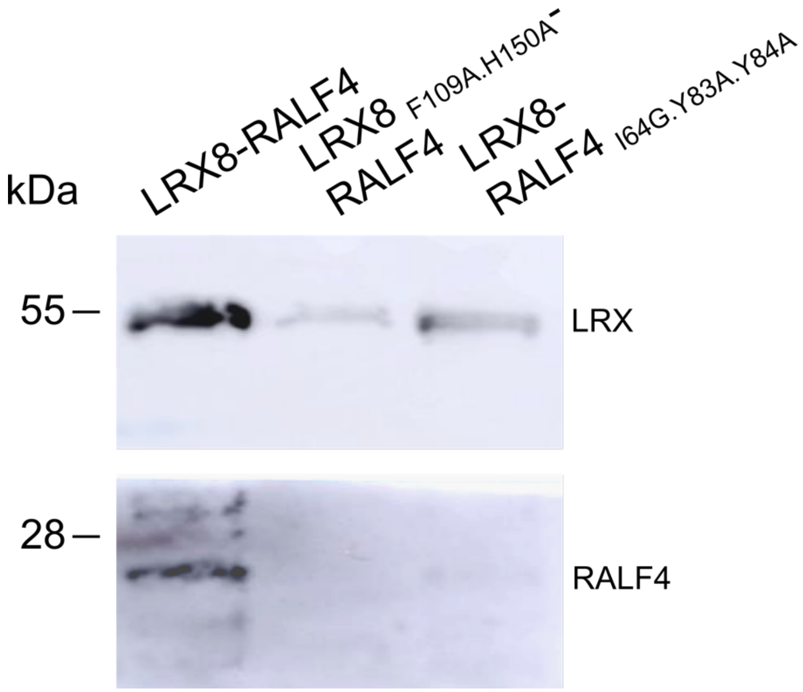

*In vitro* validation of the LRX8-RALF4 complex. Western blot from insect cell co-expression cultures of LRX8-RALF4, $LRX8_{F109A.H150A}$-RALF4 (LRX peptide binding-pocket mutant), and LRX8 – $RALF4_{I64A.Y83A.Y84A}$ (peptide binding mutant).

**Figure S10. LLG3 expression map in *Arabidopsis*.**

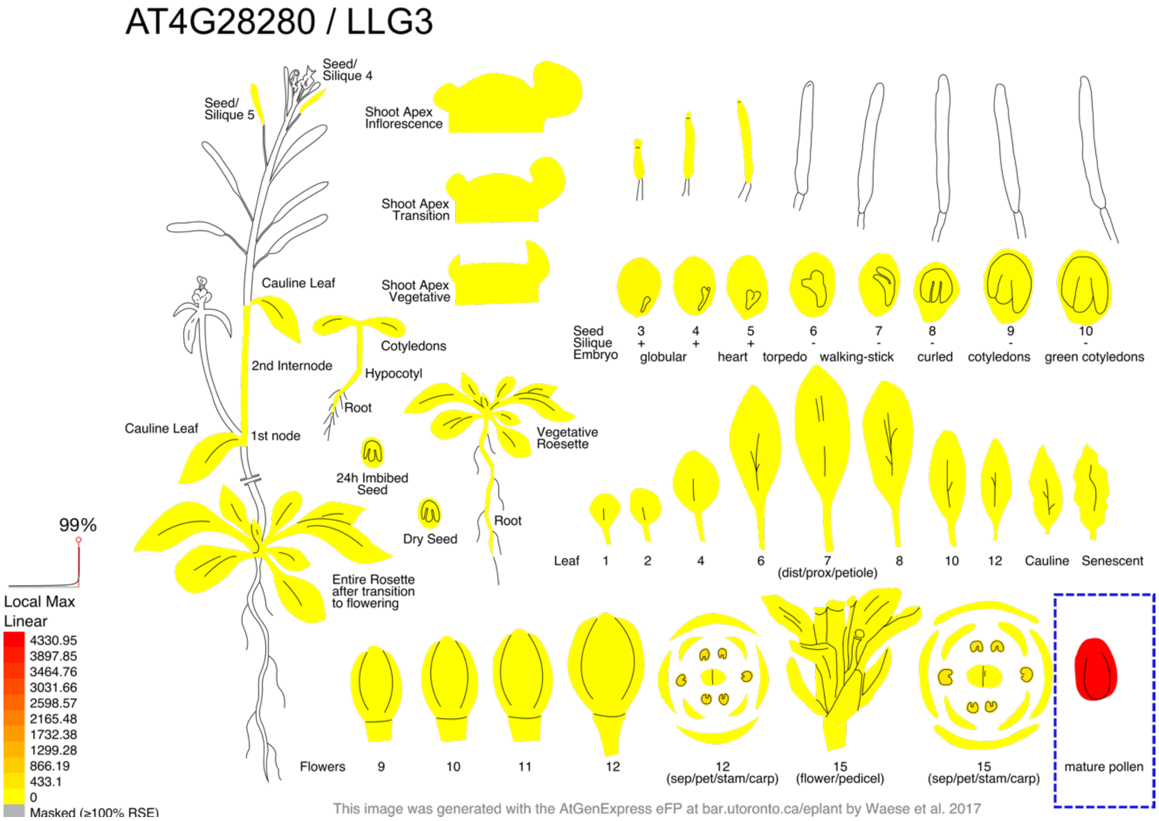

LLG3 expression in *Arabidopsis* according to public datasets available on *Arabidopsis* eFP browser (bar.utoronto.ca/efp). Relative expression levels range from low (yellow) to high (red).

**Figure S11. Purified proteins used in ITC experiments.**

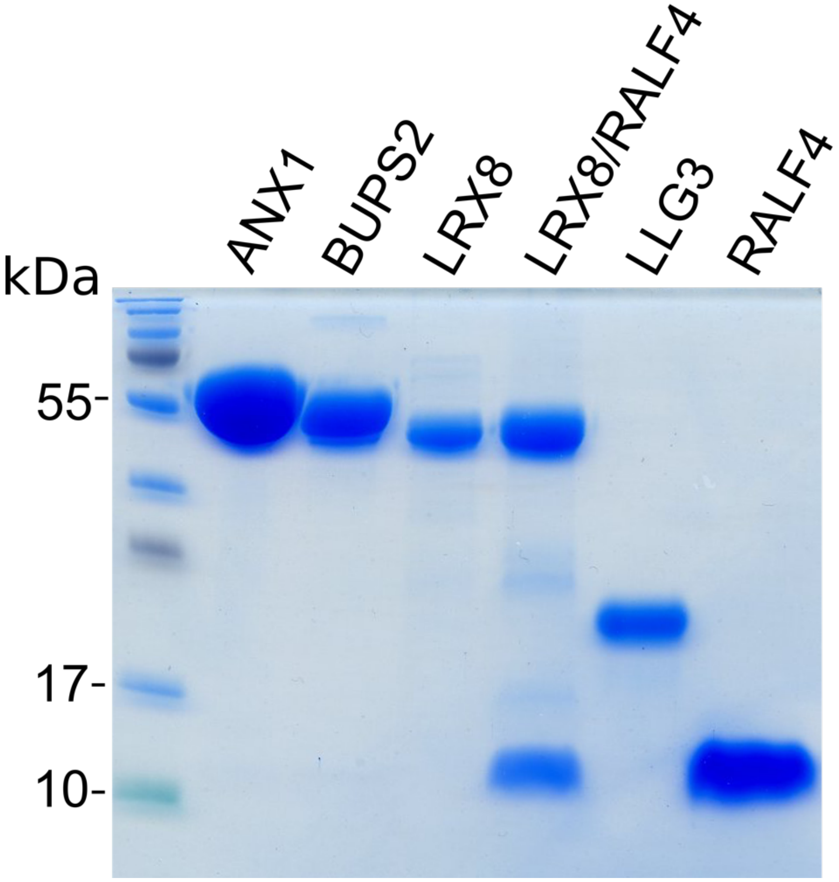

SDS-PAGE of the purified proteins used in the ITC binding matrix.

**Figure S12. Dominant active $MRI_{R240C}$ does not suppress the mutant phenotype of multiple *lrx* mutants.**

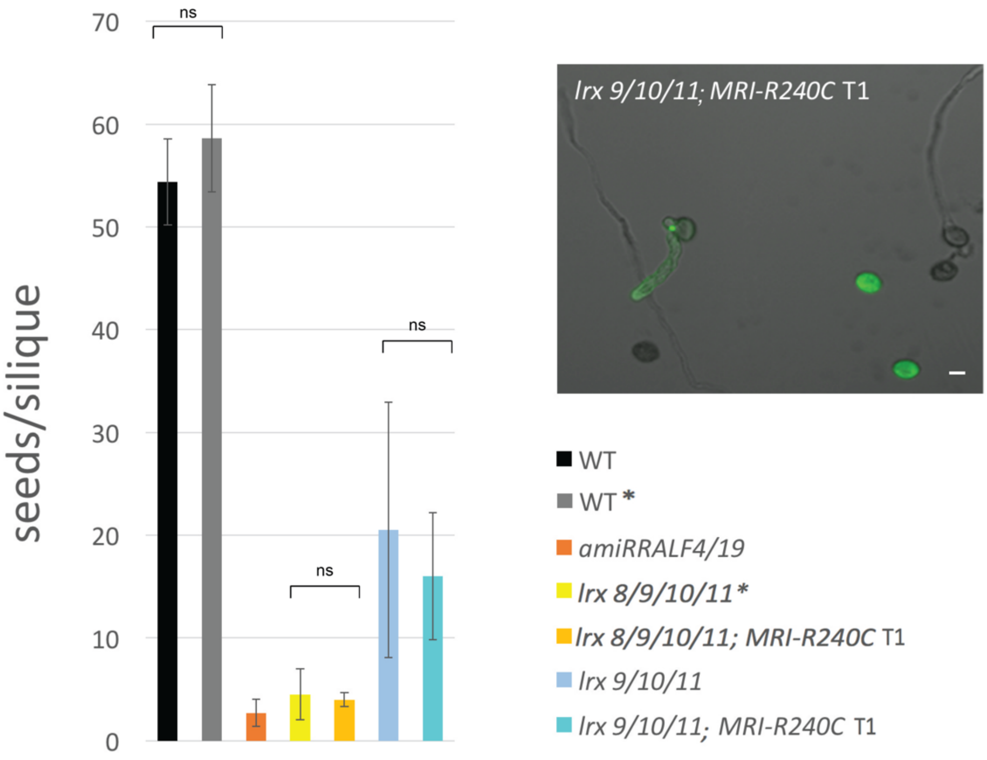

Introduction of the dominant active $MRI_{R240C}$ kinase does not suppress the pollen tube defects of *lrx8/9/10/11* quadruple and *lrx9/10/11* triple mutants, as evidenced by the strongly reduced seed set per silique. Mutants with or without the $MRI_{R240C}$ transgene (3 independent T1 lines) show no statistically significant (ns) difference in seed set. Six to eight siliques of three plants were analyzed per genotype. The plants marked by * were not grown at the same time (data from ref[5]) but since the wild-type (wt) controls showed no statistically significant difference, they are shown in the same graph. The inlet shows that although $MRI_{R240C}$-CFP was expressed in pollen grains and pollen tubes of the T1 plants, it did not suppress the *lrx9/10/11* mutant phenotype. Scale bar, 10 µm.

**Figure S13. Working model and open questions about RALF-mediated autocrine signaling during pollen tube growth.**

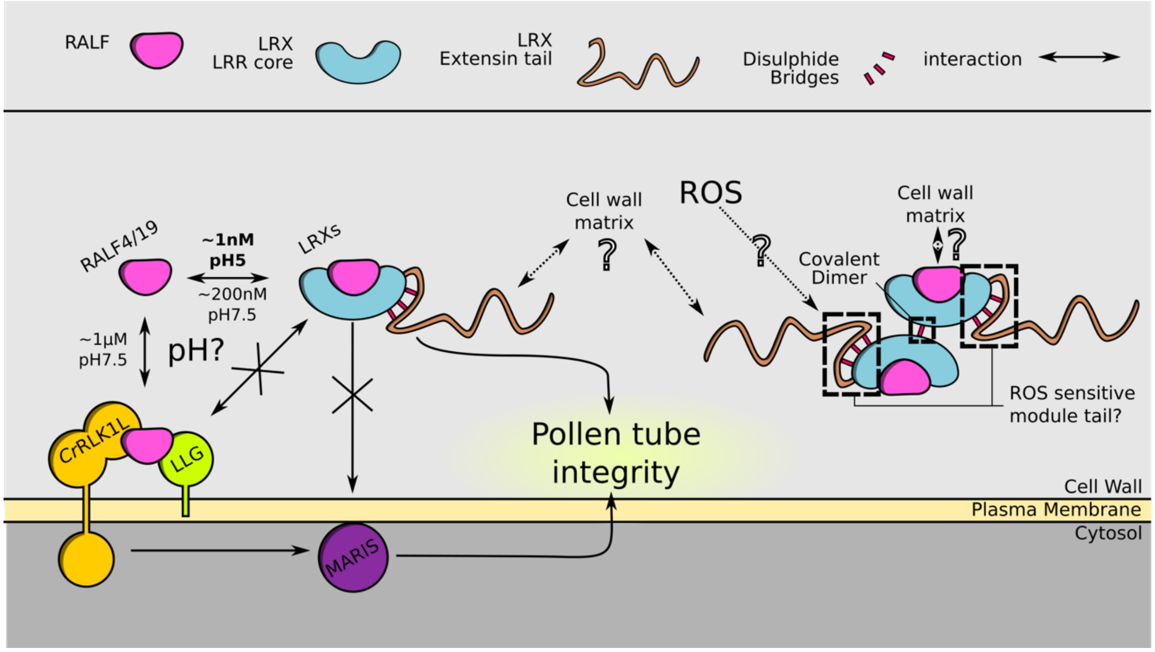

Physical interactions are depicted by double arrows, and genetic relationships by single arrows. Straight lines indicate what is known, and dotted lines speculative relationships. Affinity ranges are marked next to the interaction arrows. Legends of the different components are shown on top. Question marks indicate open questions unveiled by this study.

**Table S1. Crystallographic data-collection and refinement statistics.**

| | LRX2-RALF4 *native* | LRX8-RALF4 *native* |
|---|---|---|
| **Data collection** | | |
| Space group | *P 41* | *C 1 2 1* |
| Cell dimensions | | |
| *a*, *b*, *c* (Å) | 119.99, 119,99, 305,73 | 205.84, 114.091, 146.97 |
| $\alpha, \beta, \gamma$ (°) | 90, 90, 90 | 90, 116.25, 90 |
| Resolution (Å) | 59.99 – 3.20 (3.31 – 3.20) | 49.44 – 3.89 (4.13 – 3.89) |
| $R_{meas}$# | 0.413 (2.79) | 0.457 (2.15) |
| CC(1/2)# (%) | 99.7 (58.1) | 97.8 (34.8) |
| $I/\sigma I$# | 10.82 (1.33) | 4.44 (1.06) |
| Completeness (%)# | 99.82 (99.49) | 99.6 (99.2) |
| Redundancy# | 26.66 (27.55) | 6.7 (6.7) |
| Wilson B-factor# | 78.54 | 93.85 |
| **Refinement** | | |
| Resolution (Å) | 59.99 – 3.20 (3.31 – 3.20) | 49.44 – 3.89 (4.13 – 3.89) |
| No. reflections | 1,212,145 (184,586) | 191,042 (30,086) |
| $R_{work}/R_{free}$$ | 0.223/0.272 (0.370/0.424) | 0.281/0.329 (0.326/0.379) |
| No. atoms | | |
| Protein | 23,520 | 14,844 |
| Glycan | 773 | 226 |
| R.m.s deviations$ | | |
| Bond lengths (Å) | 0.003 | 0.003 |
| Bond angles (°) | 0.75 | 0.76 |
| Molprobity results | | |
| Ramachandran outliers (%)‡ | 0.27 | 0.32 |
| Ramachandran favored (%)‡ | 90.62 | 87.54 |
| Molprobity score‡ | 2.16 | 2.22 |
| **PDB-ID** | **6QXP** | **6QWN** |

#as defined in XDS (30), $in PHENIX (35), ‡in Molprobity (38)

**Table S2. Correspondence between amino acids of LRX2, LRX8, and LRX11.**

| Surface | LRX2 | LRX8 | LRX11 |
|---|---|---|---|
| Homodimer | C138 | C157 | C171 |
| | F68 | Y87 | F101 |
| | A114 | A133 | A147 |
| | Y116 | Y135 | Y149 |
| | R136 | R155 | R169 |
| | R160 | R179 | R193 |
| | E184 | D203 | D217 |
| LRX/RALF | F90 | F109 | V123 |
| | H131 | H150 | H164 |
| | E153 | E172 | E186 |
| | D155 | D174 | D188 |
| | D199 | D218 | D232 |
| | E248 | E266 | E280 |
| | E295 | E313 | E327 |
| | Q296 | Q314 | E328 |
| Cysteine Tail | C364 | C382 | C397 |
| | C374 | C392 | C407 |
| | C379 | C397 | C412 |
| | C324 | C253 | C267 |
| | C259 | C277 | C291 |
| | C351 | C369 | C384 |
| | I235 | I254 | I268 |
| | Q263 | E281 | E295 |
| | N326 | N344 | N358 |
| | D331 | E349 | Q363 |
| | N350 | N368 | N363 |
| | K355 | R373 | R388 |
| | R359 | K377 | R392 |
| | E363 | E381 | E396 |
| | N381 | G399 | G414 |
| N-Glycosylations | N73 | N92 | N106 |
| | N269 | N287 | N301 |
| | N320 | N338 | N352 |
| | N346 | D364 | D379 |

Table indicating the equivalent amino acids involved in the different interaction surfaces among LRX2, LRX8, and LRX11.
